## Supplementary Information for "Remodeling of gene regulatory networks underlying thermogenic stimuli-induced adipose beiging"

This PDF file includes:

- Supplementary Figure 1 to 9
- Supplementary Table 1

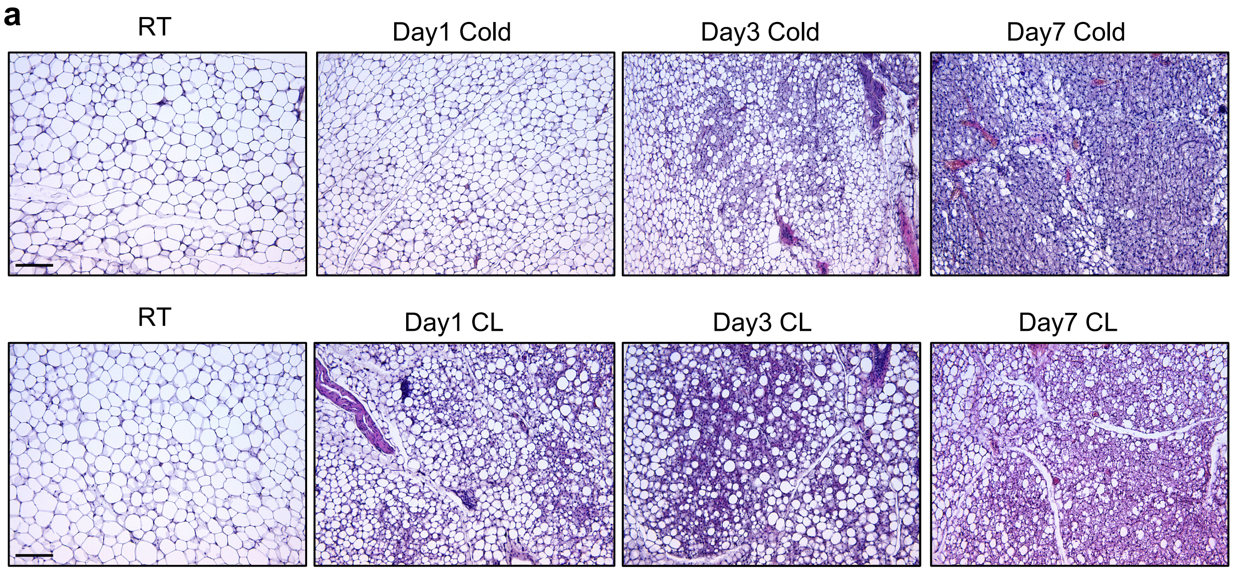


**Supplementary Fig. 1: Cold and CL lead to beige remodeling in inguinal adipose tissue.**

**a**, Representative 10X H&E-stained images of sections from iWAT depots from two-month-old male C57BL/6 mice exposed to cold (6°C) or CL-316,243 (CL; 1mg/kg/mouse/day) for 1, 3, or 7 days. Scale bar = 200 μm.

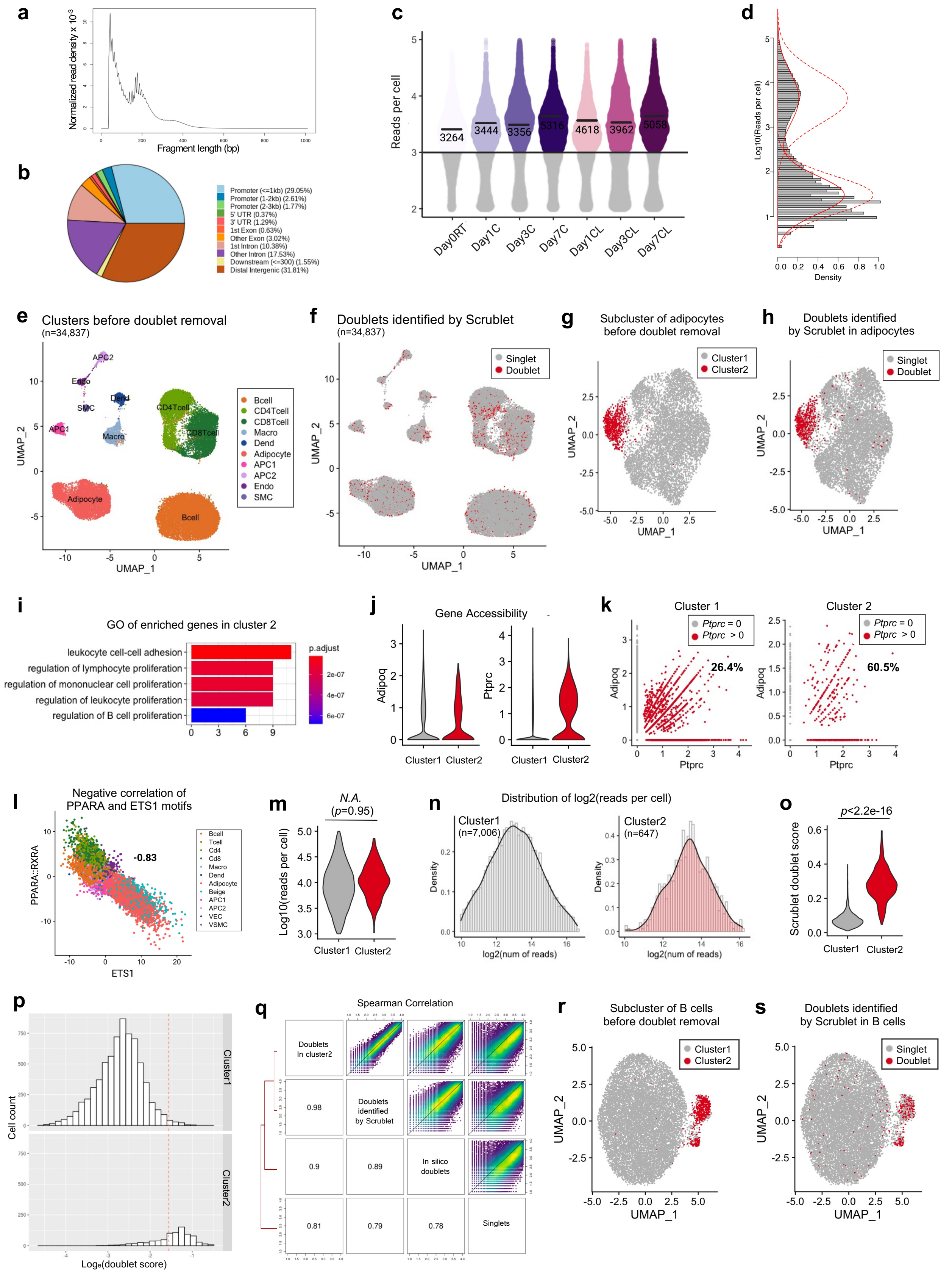
a
Normalized read density x 10⁻³
Fragment length (bp)
b
Promoter (<=1kb) (29.05%)
Promoter (1-2kb) (2.61%)
Promoter (2-3kb) (1.77%)
5' UTR (0.37%)
3' UTR (1.29%)
1st Exon (0.63%)
Other Exon (3.02%)
1st Intron (10.38%)
Other Intron (17.53%)
Downstream (<=300) (1.55%)
Distal Intergenic (31.81%)
c
Reads per cell
3264
3444
3356
5316
4618
3962
5058
Day0RT
Day1C
Day3C
Day7C
Day1CL
Day3CL
Day7CL
d
Log10(Reads per cell)
Density
e
Clusters before doublet removal
(n=34,837)
UMAP_2
UMAP_1
Bcell
CD4Tcell
CD8Tcell
Macro
Dend
Adipocyte
APC1
APC2
Endo
SMC
f
Doublets identified by Scrublet
(n=34,837)
Singlet
Doublet
g
Subcluster of adipocytes
before doublet removal
Cluster1
Cluster2
h
Doublets identified
by Scrublet in adipocytes
i
GO of enriched genes in cluster 2
leukocyte cell-cell adhesion
regulation of lymphocyte proliferation
regulation of mononuclear cell proliferation
regulation of leukocyte proliferation
regulation of B cell proliferation
p.adjust
j
Gene Accessibility
Adipoq
Ptprc
Cluster1
Cluster2
k
Cluster 1
Cluster 2
Ptprc = 0
Ptprc > 0
26.4%
60.5%
l
Negative correlation of
PPARA and ETS1 motifs
PPARA::RXRA
ETS1
-0.83
m
N.A.
(p=0.95)
Log10(reads per cell)
n
Distribution of log2(reads per cell)
Cluster1
(n=7,006)
Cluster2
(n=647)
log2(num of reads)
o
p<2.2e-16
Scrublet doublet score
p
Cell count
Logₑ(doublet score)
q
Spearman Correlation
Doublets
In cluster2
Doublets
identified
by Scrublet
In silico
doublets
Singlets
0.98
0.9
0.89
0.81
0.79
0.78
r
Subcluster of B cells
before doublet removal
s
Doublets identified
by Scrublet in B cells

**Supplementary Fig. 2: Quality control metrics and doublet identification of snATAC-seq dataset.**

**a**, Fragment size distribution plot shows peaks around 100 and 200bp, indicating enrichment of nucleosome-free and mono-nucleosome-bound fragments. **b**, Distribution of peaks on genome features shows that more than 30% of the peaks are in promoter regions, and more than half of the peaks fall into enhancer regions (distal intergenic and intronic regions). **c**, Distribution of reads per barcode for each cell from each group (n=3 for each group). The median fragment count for each group is indicated with a thick bar. The mean of log10 reads per cell of barcodes or cells passing that 1000 reads per cell cutoff is labeled. **d**, Histogram shows the distribution of reads and a bimodal distribution, indicating that a population with low reads per cell is not real cells. Red line obtained by mixture modeling separates two populations and shows inference about where the cutoff ("split") is. **e**, UMAP plot of 34,837 cells before removing doublets. Cells are colored by cell types. **f**, UMAP plot of 34,837 cells colored by grey (singlet) or red (doublet) using Scrublet[1]. **g**, UMAP plot of adipocytes (7,653 cells) before removing doublets. **h**, UMAP plot of adipocytes colored by grey (singlet) or red (doublet) using Scrublet. **i**, GO analysis of enriched genes in cluster 2. **j**, Normalized gene accessibility for adipocyte marker gene (*Adipoq*) and immune cell marker gene (*Ptprc*). **k**, Scatter plot of gene accessibility for *Adipoq* and *Ptprc*. Each dot is a cell. Red dots indicate the cells with >0 *Ptprc* accessibility. **l**, Scatter plot shows a negative correlation (-0.83 Pearson correlation) between adipocyte motif (PPARA::RXRA) and immune cell motif (ETS1). Dots are individual cells colored by cell types. **m**, Violin plot shows no statistically significant differences in reads per cell between cluster 1 and cluster 2. Welch's two sample t-test was performed. **n**, Histogram of reads per cell distribution in cluster 1 and cluster 2. **o**, Violin plot shows doublet scores of cluster 1 and cluster 2. Welch's two sample t-test was performed. **p**, Histogram of doublet score in cluster 1 and cluster 2. Red line is the threshold. **q**, Heatmap showing Pearson correlation of doublets in putative doublet cluster, doublets identified by Scrublet, in silico doublets (doublets simulated by random sampling from snATAC-seq dataset), and singlets (not in doublet cluster or identified as doublets by Scrublet). **r**, UMAP plot of B cells (12,555 cells) before removing doublets. **s**, UMAP plot of B cells colored by grey (singlet) or red (doublet) using Scrublet.

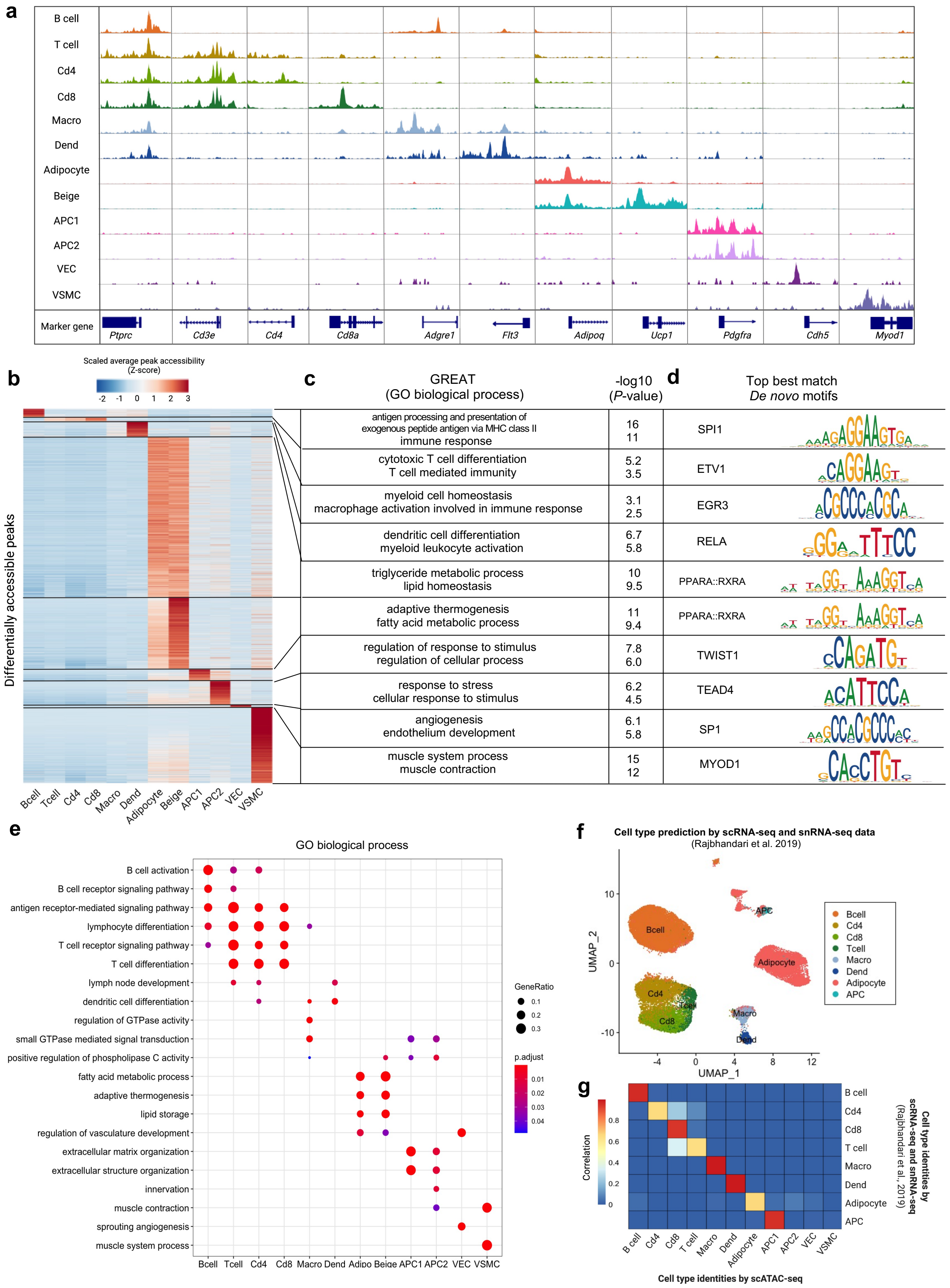
a
B cell
T cell
Cd4
Cd8
Macro
Dend
Adipocyte
Beige
APC1
APC2
VEC
VSMC
Marker gene
Ptprc
Cd3e
Cd4
Cd8a
Adgre1
Flt3
Adipoq
Ucp1
Pdgfra
Cdh5
Myod1
b
Scaled average peak accessibility (Z-score)
Differentially accessible peaks
Bcell Tcell Cd4 Cd8 Macro Dend Adipocyte Beige APC1 APC2 VEC VSMC
c
GREAT (GO biological process)
-log10 (P-value)
antigen processing and presentation of exogenous peptide antigen via MHC class II
immune response
16 11
cytotoxic T cell differentiation
T cell mediated immunity
5.2 3.5
myeloid cell homeostasis
macrophage activation involved in immune response
3.1 2.5
dendritic cell differentiation
myeloid leukocyte activation
6.7 5.8
triglyceride metabolic process
lipid homeostasis
10 9.5
adaptive thermogenesis
fatty acid metabolic process
11 9.4
regulation of response to stimulus
regulation of cellular process
7.8 6.0
response to stress
cellular response to stimulus
6.2 4.5
angiogenesis
endothelium development
6.1 5.8
muscle system process
muscle contraction
15 12
d
Top best match De novo motifs
SPI1
ETV1
EGR3
RELA
PPARA::RXRA
PPARA::RXRA
TWIST1
TEAD4
SP1
MYOD1
e
GO biological process
B cell activation
B cell receptor signaling pathway
antigen receptor-mediated signaling pathway
lymphocyte differentiation
T cell receptor signaling pathway
T cell differentiation
lymph node development
dendritic cell differentiation
regulation of GTPase activity
small GTPase mediated signal transduction
positive regulation of phospholipase C activity
fatty acid metabolic process
adaptive thermogenesis
lipid storage
regulation of vasculature development
extracellular matrix organization
extracellular structure organization
innervation
muscle contraction
sprouting angiogenesis
muscle system process
Bcell Tcell Cd4 Cd8 Macro Dend Adipo Beige APC1 APC2 VEC VSMC
GeneRatio
0.1 0.2 0.3
p.adjust
0.01 0.02 0.03 0.04
f
Cell type prediction by scRNA-seq and snRNA-seq data
(Rajbhandari et al. 2019)
UMAP_2
UMAP_1
Bcell
Cd4
Cd8
Tcell
Macro
Dend
Adipocyte
APC
g
Correlation
Cell type identities by scRNA-seq and snRNA-seq (Rajbhandari et al. 2019)
B cell
Cd4
Cd8
T cell
Macro
Dend
Adipocyte
APC
Cell type identities by scATAC-seq
B cell Cd4 Cd8 T cell Macro Dend Adipocyte APC1 APC2 VEC VSMC

**Supplementary Fig. 3: Cell type identification of mouse inguinal adipose tissue.**

**a**, Aggregated snATAC-seq accessibility profiles of the promoters for cell-type marker genes. **b**, Differentially accessible peaks (> 1.15 log2 fold-change) in each cell type. **c**, Biological function of differentially accessible regions enriched in each cell type using GREAT analysis[2] and -log10 p-value of each term. **d**, Transcription factor motif enriched in highly accessible peaks in each cell type. **e**, GO analysis using the top 50 differentially accessible genes in each cell type. **f**, UMAP plot of cells annotated by predicted cell types from scRNA-seq and snRNA-seq data[3]. Endothelial cells and smooth muscle cells were not included in scRNA-seq data. **g**, Heatmap summarizing the accuracy of cell type identification measured by Pearson correlation between clusters identified in snATAC-seq and scRNA-seq[3]. Macro; Macrophage, Dend; Dendritic cell, APC1; Adipocyte progenitor cell 1, APC2; Adipocyte progenitor cell 2, VEC; Vascular endothelial cell, VSMC; Vascular smooth muscle cell

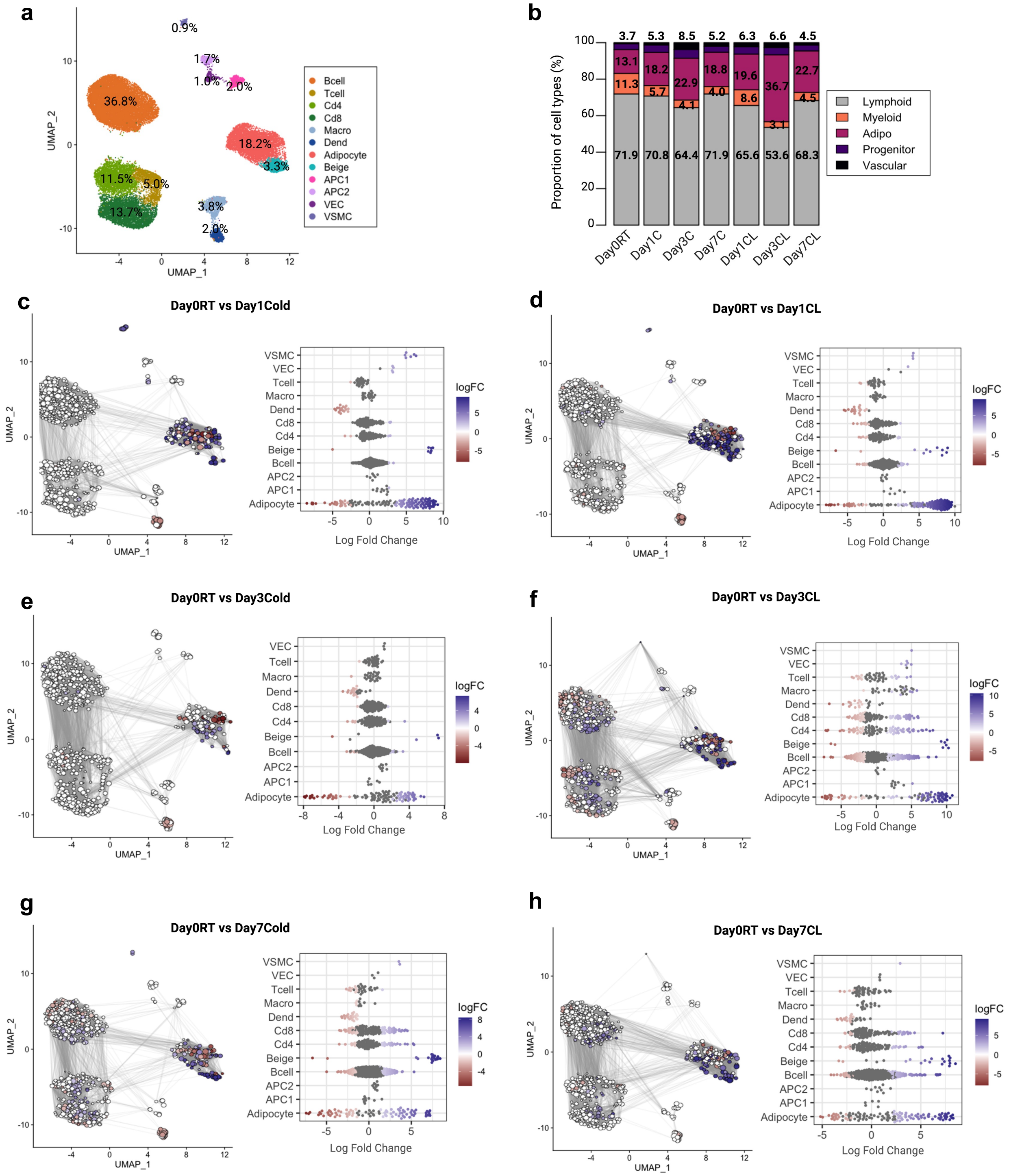


**Supplementary Fig. 4: Cell type distribution and differential cell abundance after cold exposure and CL treatment.**

**a**, UMAP plot of 32,552 cells from all groups. The percentage of cell types is labeled. **b**, Proportion of major cell types in each group. **c**-**h**, Milo analysis[4] of cell neighborhood abundance changes in Day 0 control *vs*.

Day 1, 3, or 7 of cold (**c**,**e**,**g**) or CL (**d**,**f**,**h**). In UMAP, size of points indicates the number of cells in a neighborhood; lines represent the number of cells shared between adjacent neighborhoods. Points are neighborhoods (Nhood), colored by the log fold differences at the two time points (FDR 10%). Beeswarm plots show the distribution of the log-fold in defined clusters.

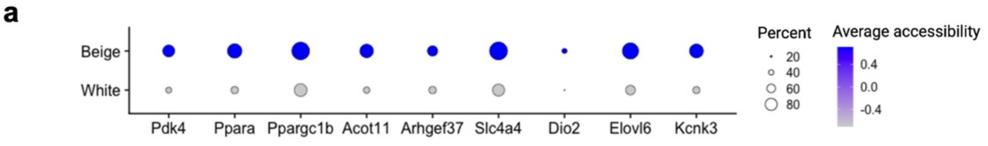


**b**

| | H3K4me1 | | | H3K27ac | | | "poised" H3K4me1 in the absence of H3K27ac | | |
|---|---|---|---|---|---|---|---|---|---|
| | cold beige | warm beige | warm white | cold beige | warm beige | warm white | cold beige | warm beige | warm white |
| white | 371 | 464 | 457 | 1600 | 1641 | 1867 | 1466 | 1344 | 1581 |
| beige | 1626 | 1654 | 1778 | 6424 | 5989 | 4814 | 5657 | 4594 | 3742 |

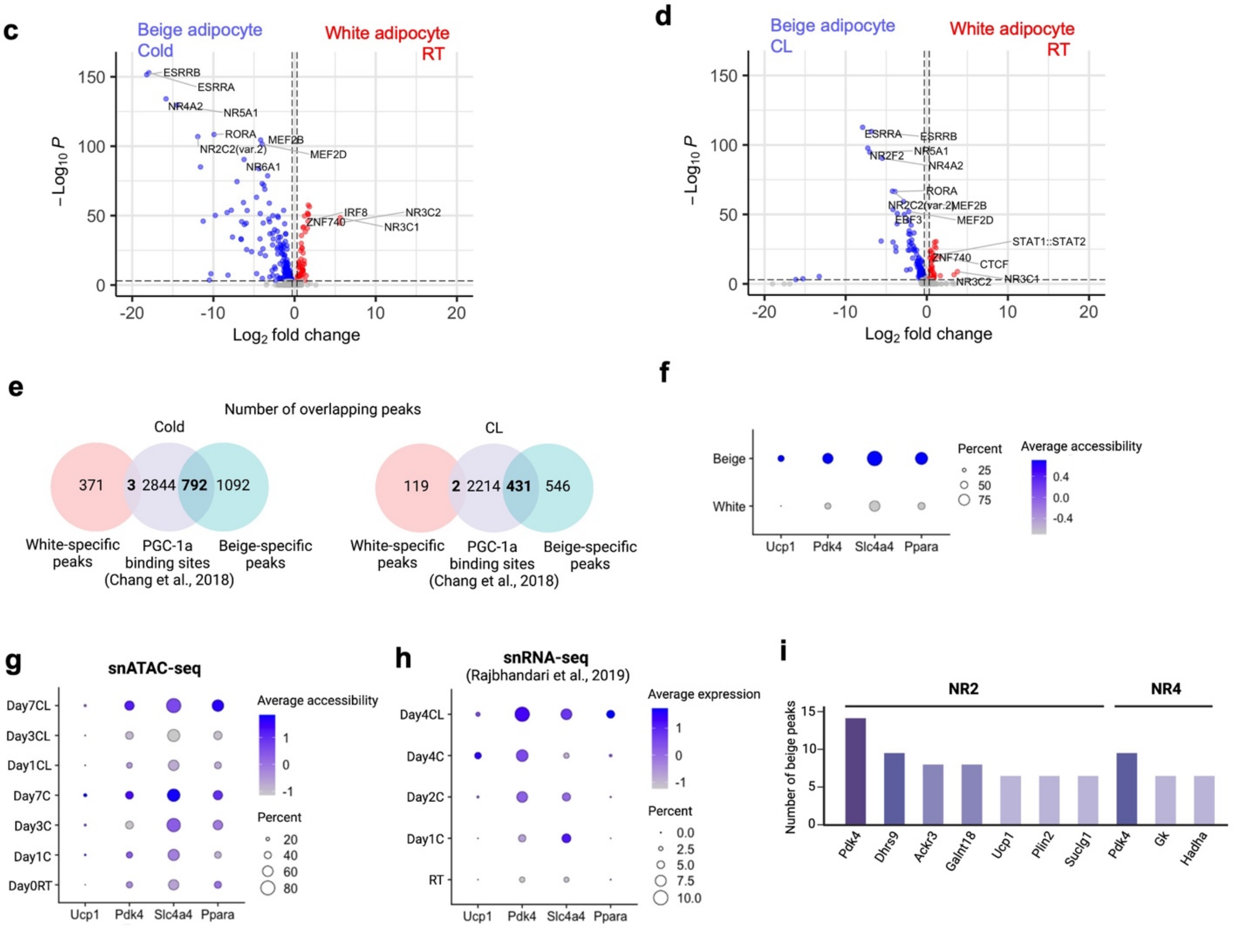


**Supplementary Fig. 5: Coordinating regulation of transcription factors and epigenetic modifiers in white and beige adipocytes.**

**a**, Dot plot showing normalized accessibility of genes with more than three beige-specific peaks closest to them. Dot size depicts the percent of cells having accessibility of a given gene. **b**, The number of overlaps between white-specific or beige-specific peaks from snATAC-seq data and histone marked peaks from H3Kme1 and H3K27ac ChIP-seq data[5]. White specific peaks tend to overlap with peaks enriched in warm white, while beige specific peaks tend to overlap with peaks enriched in cold beige. **c**,**d**, Volcano plot displaying differential motifs enrichment between white adipocytes before any intervention and beige adipocytes after cold exposure (**c**) or CL treatment (**d**). Motifs with adjusted *p*-value < $10^{-3}$ & Abs(log2 fold-change) > 0.5 are colored. **e**, Venn diagram showing the number of overlaps between white-specific or beige-specific peaks after cold exposure or CL treatment and PGC-1α ChIP-seq binding sites[6]. **f**, Normalized accessibilities of the genes having more than three nearby beige-specific peaks that are overlapping with PGC-1α binding sites. **g,h**, Dot plots displaying normalized snATAC-seq accessibility (**g**) and snRNA-seq expression level[3] (**h**) of the genes having more than three nearby beige-specific peaks overlapping with PGC-1α binding sites. **i**, The number of overlapping beige-specific peaks with NR2 and NR4 motifs from *cisbp* database[7]. Genes are sorted by the number of overlapping peaks near them.

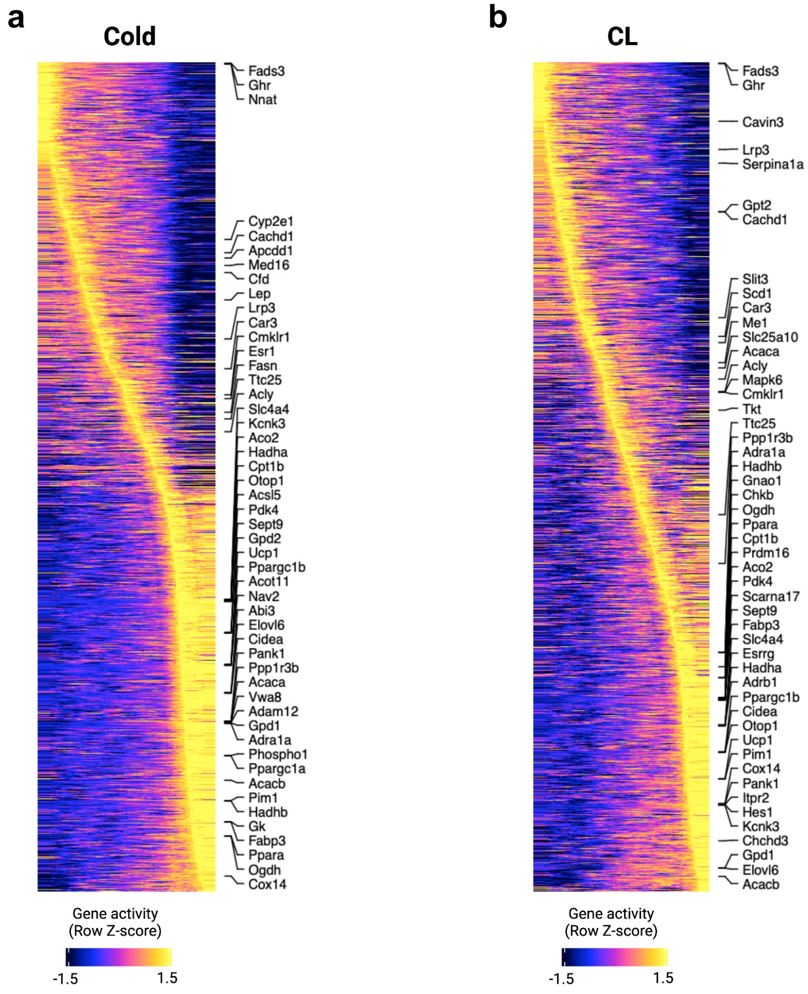


**Supplementary Fig. 6: Gene accessibility changes along with adipocyte pseudotime trajectory.**

**a**,**b**, Heatmaps display changes in gene accessibility along adipocyte pseudotime trajectory for cold (**a**) and CL (**b**). Top variable genes are labeled on the right side of the heatmap.

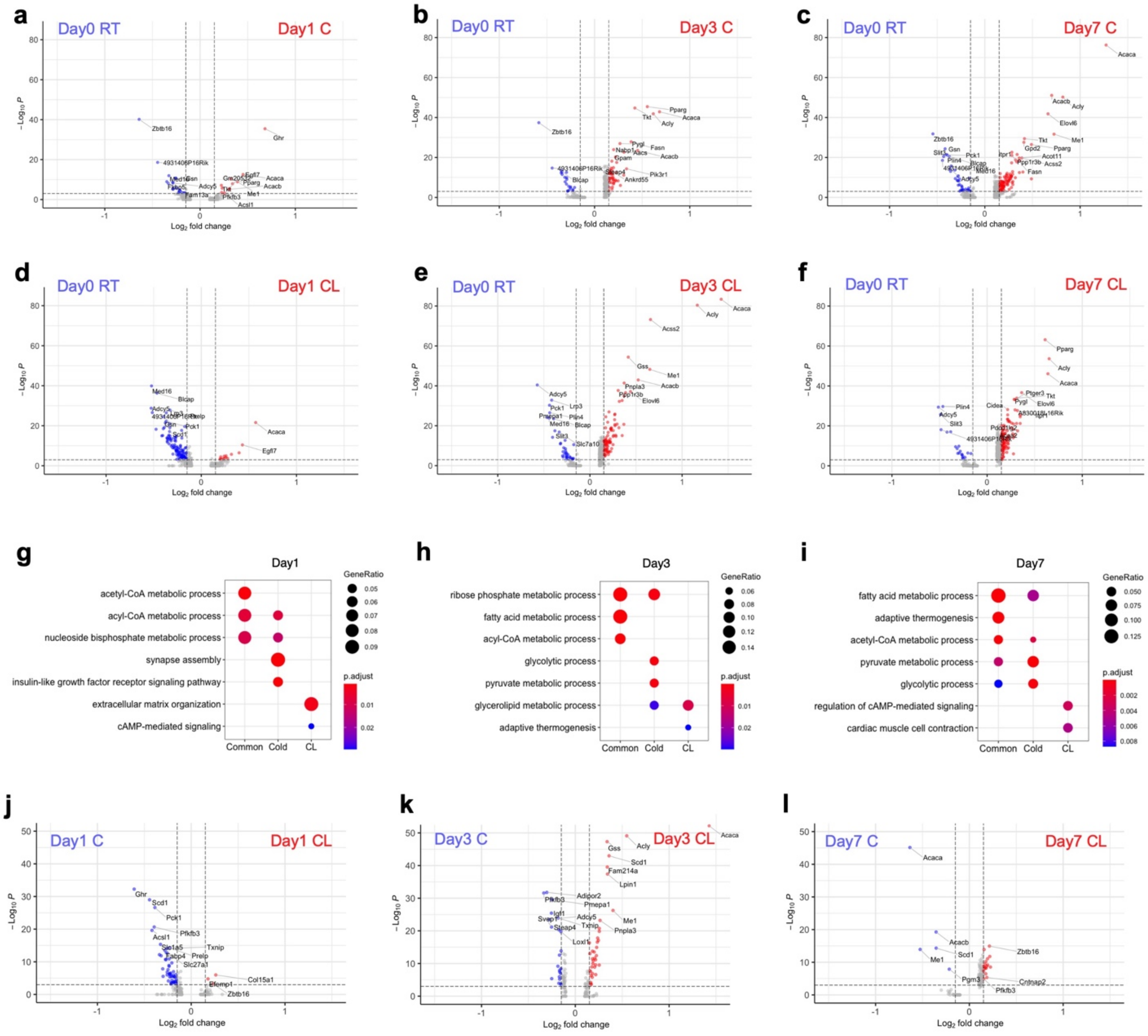


**Supplementary Fig. 7: Differentially accessible genes after cold exposure or CL treatment in adipocytes.**

**a**-**f**, Volcano plots of differentially accessible genes between day 0 RT and cold (**a**,**b**,**c**) or CL (**d**,**e**,**f**) at each time point in mature adipocytes (both white and beige adipocyte). Genes with adjusted *p*-value < 0.001 & Abs(log10 fold-change) > 0.15 are colored and labeled. **g**-**i**, GO analysis of commonly more accessible genes after cold and CL treatment (common) and uniquely more accessible in cold or CL at each time point. **j**-**l**, Volcano plots of the differentially accessible genes between cold and CL at day 1 (**j**), day 3 (**k**), and day 7 (**l**) in mature adipocytes (both white and beige). Genes with adjusted *p*-value < 0.001 & Abs(log2 fold-change) > 0.15 are colored and labeled.

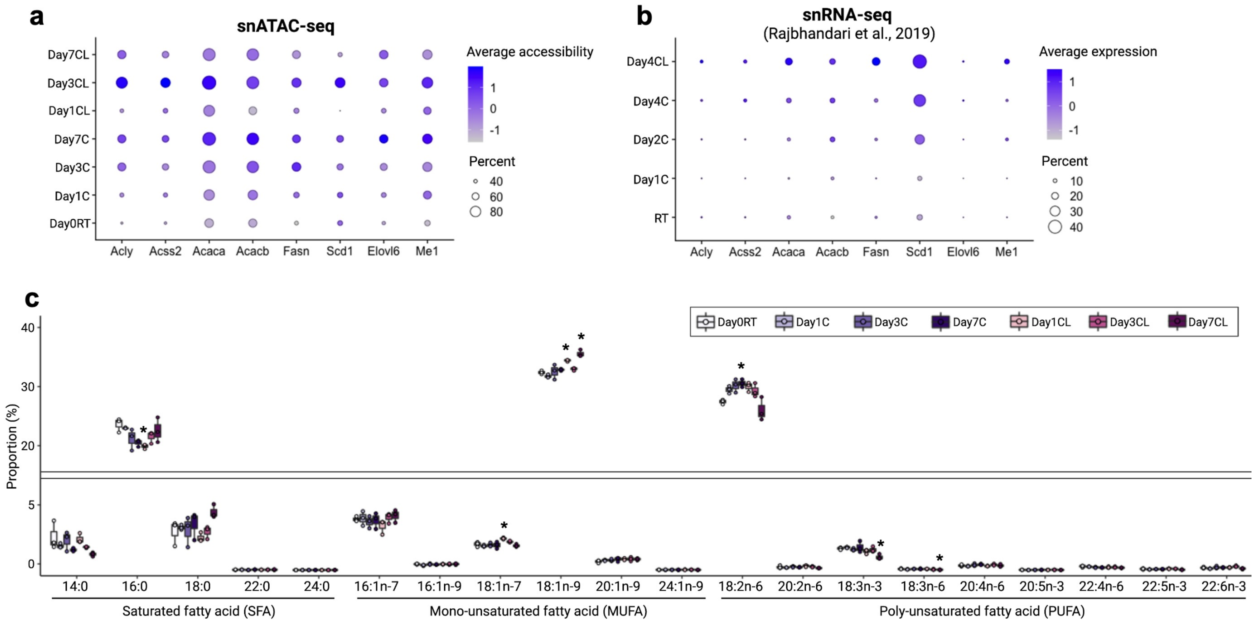


**Supplementary Fig. 8: Changes in accessibility and expression of genes involved in lipogenesis and proportion of lipid species after cold and CL treatment.**

**a**,**b**, Dot plots displaying normalized snATAC-seq accessibility (**a**) and snRNA-seq expression level[3] (**b**) of the lipogenic genes. **c**, Relative proportion of each lipid class by lipidomics analysis (n=3 per group). Color of bar indicates group. *Adjusted *p*-value<0.05, ANOVA multiple comparisons test with Bonferroni's post-hoc test was performed.

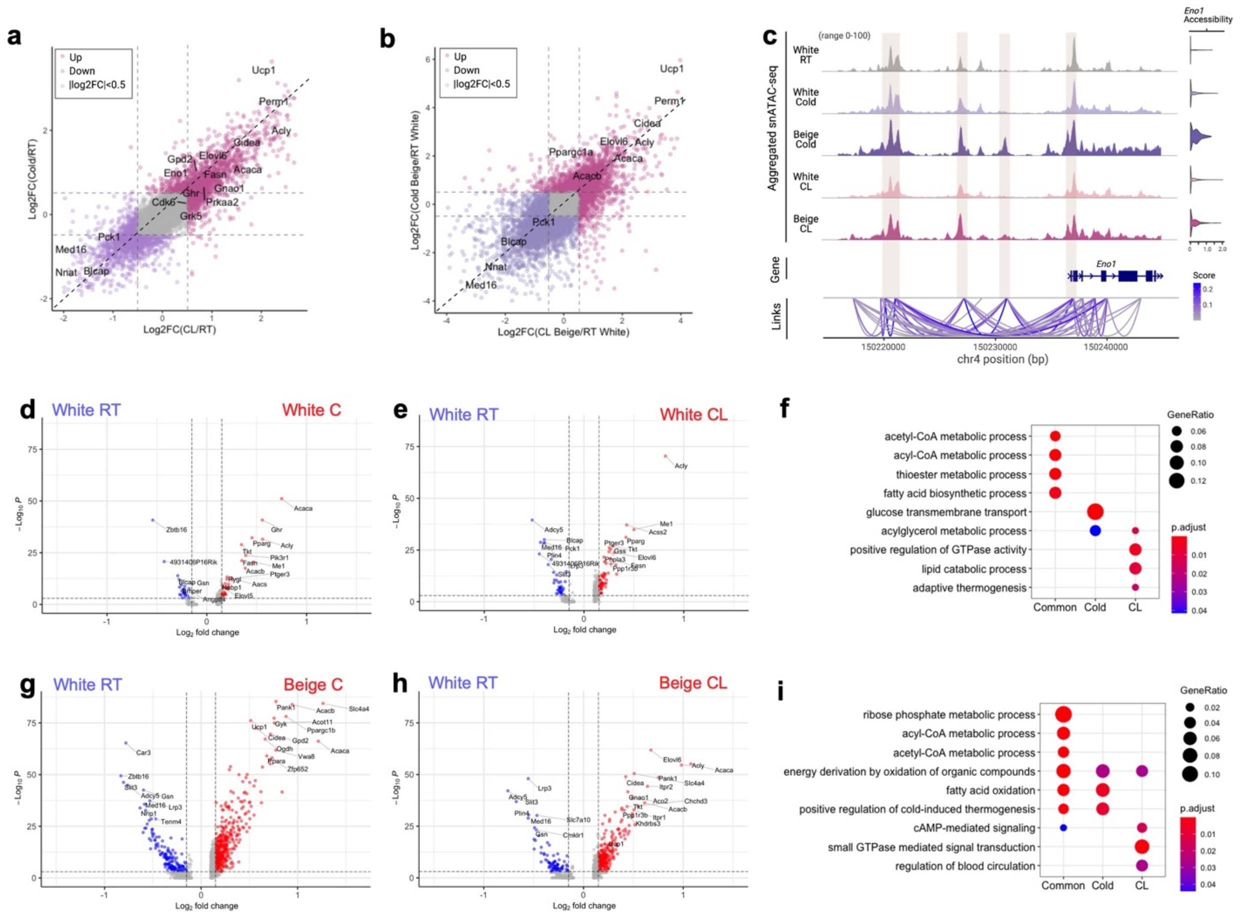


**Supplementary Fig. 9: Differentially accessible genes after cold exposure and CL treatment in white and beige adipocytes.**

**a**, Scatter plot of the log2 fold-change of genes after cold exposure and CL treatment. The genes with >0.5 log2 fold changes are colored. **b**, Scatter plot of the log2 fold-change of genes in cold-induced and CL-induced beige adipocytes compared to white adipocytes at RT. The genes with >0.5 log2 fold changes are colored. **c**, Genome tracks showing the aggregate snATAC-seq profiles of *Eno1* upstream in adipocytes. Links show *cis*-coaccessibility network with multiple connections between peaks around *Eno1*. Coaccessibility score > 0.2 regions are highlighted by brown on genome and gene track. **d,e**, Volcano plots of the differentially accessible genes between white adipocytes at day 0 RT and after cold exposure (**d**) or CL treatment (**e**) from all three time points. Genes with adjusted *p*-value < 0.0001 & Abs(log2 fold-change) > 0.15 are colored. **f**, GO analysis of commonly more accessible genes after cold exposure and CL treatment (common) and uniquely more accessible after cold exposure or CL treatment in white adipocytes. **g**,**h**, Volcano plots of the differentially accessible genes between white adipocytes at day 0 RT and beige adipocytes after cold exposure (**g**) or CL treatment (**h**) from all three time points. Genes with adjusted *p*-value < 0.0001 & Abs(log2 fold-change) > 0.15 are colored. **i**, GO analysis of commonly more accessible genes after cold exposure and CL treatment (common) and uniquely more accessible after cold exposure or CL treatment in beige adipocytes.

**Supplementary Table 1. Oligonucleotide sequences.**

| Oligo | Sequence (5'-3') |
|---|---|
| Tn5ME_Rev | 5Phos/CTGTCTCTTATACACATCT |
| P5_ME_1 | TCGTCGGCAGCGTCTCCACGCTATAGCCTGCGATCGAGGACGGCAGATGTGTATAAGAGACAG |
| P5_ME_2 | TCGTCGGCAGCGTCTCCACGCATAGAGGCGCGATCGAGGACGGCAGATGTGTATAAGAGACAG |
| P5_ME_3 | TCGTCGGCAGCGTCTCCACGCCCTATCCTGCGATCGAGGACGGCAGATGTGTATAAGAGACAG |
| P5_ME_4 | TCGTCGGCAGCGTCTCCACGCGGCTCTGAGCGATCGAGGACGGCAGATGTGTATAAGAGACAG |
| P5_ME_5 | TCGTCGGCAGCGTCTCCACGCAGGCGAAGGCGATCGAGGACGGCAGATGTGTATAAGAGACAG |
| P5_ME_6 | TCGTCGGCAGCGTCTCCACGCTAATCTTAGCGATCGAGGACGGCAGATGTGTATAAGAGACAG |
| P5_ME_7 | TCGTCGGCAGCGTCTCCACGCCAGGACGTGCGATCGAGGACGGCAGATGTGTATAAGAGACAG |
| P5_ME_8 | TCGTCGGCAGCGTCTCCACGCGTACTGACGCGATCGAGGACGGCAGATGTGTATAAGAGACAG |
| P7_ME_1 | GTCTCGTGGGCTCGGCTGTCCCTGTCCCGAGTAATCACCGTCTCCGCCTCAGATGTGTATAAGAGACAG |
| P7_ME_2 | GTCTCGTGGGCTCGGCTGTCCCTGTCCTCTCCGGACACCGTCTCCGCCTCAGATGTGTATAAGAGACAG |
| P7_ME_3 | GTCTCGTGGGCTCGGCTGTCCCTGTCCAATGAGCGCACCGTCTCCGCCTCAGATGTGTATAAGAGACAG |
| P7_ME_4 | GTCTCGTGGGCTCGGCTGTCCCTGTCCGGAATCTCCACCGTCTCCGCCTCAGATGTGTATAAGAGACAG |
| P7_ME_5 | GTCTCGTGGGCTCGGCTGTCCCTGTCCTTCTGAATCACCGTCTCCGCCTCAGATGTGTATAAGAGACAG |
| P7_ME_6 | GTCTCGTGGGCTCGGCTGTCCCTGTCCACGAATTCCACCGTCTCCGCCTCAGATGTGTATAAGAGACAG |
| P7_ME_7 | GTCTCGTGGGCTCGGCTGTCCCTGTCCAGCTTCAGCACCGTCTCCGCCTCAGATGTGTATAAGAGACAG |
| P7_ME_8 | GTCTCGTGGGCTCGGCTGTCCCTGTCCGCGCATTACACCGTCTCCGCCTCAGATGTGTATAAGAGACAG |
| P7_ME_9 | GTCTCGTGGGCTCGGCTGTCCCTGTCCCATAGCCGCACCGTCTCCGCCTCAGATGTGTATAAGAGACAG |
| P7_ME_10 | GTCTCGTGGGCTCGGCTGTCCCTGTCCTTCGCGGACACCGTCTCCGCCTCAGATGTGTATAAGAGACAG |
| P7_ME_11 | GTCTCGTGGGCTCGGCTGTCCCTGTCCGCGCGAGACACCGTCTCCGCCTCAGATGTGTATAAGAGACAG |
| P7_ME_12 | GTCTCGTGGGCTCGGCTGTCCCTGTCCCTATCGCTCACCGTCTCCGCCTCAGATGTGTATAAGAGACAG |
| I5_1 | AATGATACGGCGACCACCGAGATCTACACCTCTCTATTCGTCGGCAGCGTC |
| I5_2 | AATGATACGGCGACCACCGAGATCTACACTATCCTCTTCGTCGGCAGCGTC |
| I5_3 | AATGATACGGCGACCACCGAGATCTACACGTAAGGAGTCGTCGGCAGCGTC |
| I5_4 | AATGATACGGCGACCACCGAGATCTACACACTGCATATCGTCGGCAGCGTC |
| I5_5 | AATGATACGGCGACCACCGAGATCTACACAAGGAGTATCGTCGGCAGCGTC |
| I5_6 | AATGATACGGCGACCACCGAGATCTACACCTAAGCCTTCGTCGGCAGCGTC |
| I5_7 | AATGATACGGCGACCACCGAGATCTACACCGTCTAATTCGTCGGCAGCGTC |
| I5_8 | AATGATACGGCGACCACCGAGATCTACACTCTCTCCGTCGTCGGCAGCGTC |
| I5_9 | AATGATACGGCGACCACCGAGATCTACACTCGACTAGTCGTCGGCAGCGTC |
| I5_10 | AATGATACGGCGACCACCGAGATCTACACTTCTAGCTTCGTCGGCAGCGTC |
| I5_11 | AATGATACGGCGACCACCGAGATCTACACCCTAGAGTTCGTCGGCAGCGTC |
| I5_12 | AATGATACGGCGACCACCGAGATCTACACGCGTAAGATCGTCGGCAGCGTC |
| I5_13 | AATGATACGGCGACCACCGAGATCTACACAAGGCTATTCGTCGGCAGCGTC |
| I5_14 | AATGATACGGCGACCACCGAGATCTACACGAGCCTTATCGTCGGCAGCGTC |
| I5_15 | AATGATACGGCGACCACCGAGATCTACACTTATGCGATCGTCGGCAGCGTC |
| I5_16 | AATGATACGGCGACCACCGAGATCTACACATCTGAGTTCGTCGGCAGCGTC |
| I5_17 | AATGATACGGCGACCACCGAGATCTACACGGATACTATCGTCGGCAGCGTC |
| I5_18 | AATGATACGGCGACCACCGAGATCTACACTAAGATCCTCGTCGGCAGCGTC |
| I5_19 | AATGATACGGCGACCACCGAGATCTACACAAGAGATGTCGTCGGCAGCGTC |
| I5_20 | AATGATACGGCGACCACCGAGATCTACACAATGACGTTCGTCGGCAGCGTC |
| I5_21 | AATGATACGGCGACCACCGAGATCTACACGAAGTATGTCGTCGGCAGCGTC |
| I5_22 | AATGATACGGCGACCACCGAGATCTACACATAGCCTTTCGTCGGCAGCGTC |
| I5_23 | AATGATACGGCGACCACCGAGATCTACACTTGGAAGTTCGTCGGCAGCGTC |
| I5_24 | AATGATACGGCGACCACCGAGATCTACACATTCGTTGTCGTCGGCAGCGTC |
| I5_25 | AATGATACGGCGACCACCGAGATCTACACAGGATAACTCGTCGGCAGCGTC |
| I5_26 | AATGATACGGCGACCACCGAGATCTACACTTCATCCATCGTCGGCAGCGTC |
| I5_27 | AATGATACGGCGACCACCGAGATCTACACAACGAACGTCGTCGGCAGCGTC |
| I5_28 | AATGATACGGCGACCACCGAGATCTACACTGCCTTACTCGTCGGCAGCGTC |
| I5_29 | AATGATACGGCGACCACCGAGATCTACACCGAATTCCTCGTCGGCAGCGTC |
| I5_30 | AATGATACGGCGACCACCGAGATCTACACGGTTAGACTCGTCGGCAGCGTC |
| I5_31 | AATGATACGGCGACCACCGAGATCTACACTCCGGTAATCGTCGGCAGCGTC |
| I5_32 | AATGATACGGCGACCACCGAGATCTACACTTACGACCTCGTCGGCAGCGTC |
| I7_1 | CAAGCAGAAGACGGCATACGAGATTCGCCTTAGTCTCGTGGGCTCGG |
| I7_2 | CAAGCAGAAGACGGCATACGAGATCTAGTACGGTCTCGTGGGCTCGG |
| I7_3 | CAAGCAGAAGACGGCATACGAGATTTCTGCCTGTCTCGTGGGCTCGG |
| I7_4 | CAAGCAGAAGACGGCATACGAGATGCTCAGGAGTCTCGTGGGCTCGG |
| I7_5 | CAAGCAGAAGACGGCATACGAGATAGGAGTCCGTCTCGTGGGCTCGG |

| | |
|---|---|
| I7_6 | CAAGCAGAAGACGGCATACGAGATCATGCCTAGTCTCGTGGGCTCGG |
| I7_7 | CAAGCAGAAGACGGCATACGAGATGTAGAGAGGTCTCGTGGGCTCGG |
| I7_8 | CAAGCAGAAGACGGCATACGAGATCAGCCTCGGTCTCGTGGGCTCGG |
| I7_9 | CAAGCAGAAGACGGCATACGAGATTGCCTCTTGTCTCGTGGGCTCGG |
| I7_10 | CAAGCAGAAGACGGCATACGAGATTCCTCTACGTCTCGTGGGCTCGG |
| I7_11 | CAAGCAGAAGACGGCATACGAGATTCATGAGCGTCTCGTGGGCTCGG |
| I7_12 | CAAGCAGAAGACGGCATACGAGATCCTGAGATGTCTCGTGGGCTCGG |
| I7_13 | CAAGCAGAAGACGGCATACGAGATTAGCGAGTGTCTCGTGGGCTCGG |
| I7_14 | CAAGCAGAAGACGGCATACGAGATGTAGCTCCGTCTCGTGGGCTCGG |
| I7_15 | CAAGCAGAAGACGGCATACGAGATTACTACGCGTCTCGTGGGCTCGG |
| I7_16 | CAAGCAGAAGACGGCATACGAGATGCAGCGTAGTCTCGTGGGCTCGG |
| I7_17 | CAAGCAGAAGACGGCATACGAGATCTGCGCATGTCTCGTGGGCTCGG |
| I7_18 | CAAGCAGAAGACGGCATACGAGATGAGCGCTAGTCTCGTGGGCTCGG |
| I7_19 | CAAGCAGAAGACGGCATACGAGATCGCTCAGTGTCTCGTGGGCTCGG |
| I7_20 | CAAGCAGAAGACGGCATACGAGATGTCTTAGGGTCTCGTGGGCTCGG |
| I7_21 | CAAGCAGAAGACGGCATACGAGATACTGATCGGTCTCGTGGGCTCGG |
| I7_22 | CAAGCAGAAGACGGCATACGAGATTAGCTGCAGTCTCGTGGGCTCGG |
| I7_23 | CAAGCAGAAGACGGCATACGAGATGACGTCGAGTCTCGTGGGCTCGG |
| I7_24 | CAAGCAGAAGACGGCATACGAGATTACCAGAGGTCTCGTGGGCTCGG |
| I7_25 | CAAGCAGAAGACGGCATACGAGATGGATGGAAGTCTCGTGGGCTCGG |
| I7_26 | CAAGCAGAAGACGGCATACGAGATATTGAGGCGTCTCGTGGGCTCGG |
| I7_27 | CAAGCAGAAGACGGCATACGAGATCGGATAGAGTCTCGTGGGCTCGG |
| I7_28 | CAAGCAGAAGACGGCATACGAGATTGGTAGACGTCTCGTGGGCTCGG |
| I7_29 | CAAGCAGAAGACGGCATACGAGATACCTGGTTGTCTCGTGGGCTCGG |
| I7_30 | CAAGCAGAAGACGGCATACGAGATCAGTTCTGGTCTCGTGGGCTCGG |
| I7_31 | CAAGCAGAAGACGGCATACGAGATTCGAACGTGTCTCGTGGGCTCGG |
| I7_32 | CAAGCAGAAGACGGCATACGAGATCGTTGCTTGTCTCGTGGGCTCGG |
| I7_33 | CAAGCAGAAGACGGCATACGAGATTACCGTTCGTCTCGTGGGCTCGG |
| I7_34 | CAAGCAGAAGACGGCATACGAGATTAGGTTGCGTCTCGTGGGCTCGG |
| I7_35 | CAAGCAGAAGACGGCATACGAGATGAGGCTAAGTCTCGTGGGCTCGG |
| I7_36 | CAAGCAGAAGACGGCATACGAGATCGACCATAGTCTCGTGGGCTCGG |
| I7_37 | CAAGCAGAAGACGGCATACGAGATAGGCAGTAGTCTCGTGGGCTCGG |
| I7_38 | CAAGCAGAAGACGGCATACGAGATATCAAGCGGTCTCGTGGGCTCGG |
| I7_39 | CAAGCAGAAGACGGCATACGAGATCATTGAAGGTCTCGTGGGCTCGG |
| I7_40 | CAAGCAGAAGACGGCATACGAGATCGACTTATGTCTCGTGGGCTCGG |
| I7_41 | CAAGCAGAAGACGGCATACGAGATTCTATACGGTCTCGTGGGCTCGG |
| I7_42 | CAAGCAGAAGACGGCATACGAGATAGCATTAGGTCTCGTGGGCTCGG |
| I7_43 | CAAGCAGAAGACGGCATACGAGATAATTGGCAGTCTCGTGGGCTCGG |
| I7_44 | CAAGCAGAAGACGGCATACGAGATAGATTCGTGTCTCGTGGGCTCGG |
| I7_45 | CAAGCAGAAGACGGCATACGAGATTTCATGACGTCTCGTGGGCTCGG |
| I7_46 | CAAGCAGAAGACGGCATACGAGATTGAACTTGGTCTCGTGGGCTCGG |
| I7_47 | CAAGCAGAAGACGGCATACGAGATATGGCATAGTCTCGTGGGCTCGG |
| I7_48 | CAAGCAGAAGACGGCATACGAGATCGTAATTCGTCTCGTGGGCTCGG |
| Read 1 Sequencing Primer | GCGATCGAGGACGGCAGATGTGTATAAGAGACAG |
| Read 2 Sequencing Primer | CACCGTCTCCGCCTCAGATGTGTATAAGAGACAG |
| Index 1 Sequencing Primer | CTGTCTCTTATACACATCTGAGGCGGAGACGGTG |
| Index 2 Sequencing Primer | CTGTCTCTTATACACATCTGCCGTCCTCGATCGC |
